# Generation and analysis of a mouse multi-tissue genome annotation atlas

**DOI:** 10.1101/2024.01.31.578267

**Authors:** Matthew Adams, Christopher Vollmers

**Affiliations:** Department of Molecular, Cellular, and Developmental Biology, University of California Santa Cruz; Department of Biomolecular Engineering, University of California Santa Cruz

## Abstract

Generating an accurate and complete genome annotation for an organism is complex because the cells within each tissue can express a unique set of transcript isoforms from a unique set of genes. A comprehensive genome annotation should contain information on what tissues express what transcript isoforms at what level. This tissue-level isoform information can then inform a wide range of research questions as well as experiment designs. Long-read sequencing technology combined with advanced full-length cDNA library preparation methods has now achieved throughput and accuracy where generating these types of annotations is achievable.

Here, we show this by generating a genome annotation of the mouse (Mus musculus). We used the nanopore-based R2C2 long-read sequencing method to generate 64 million highly accurate full length cDNA consensus reads - averaging 5.4 million reads per tissue for a dozen tissues. Using the Mandalorion tool we processed these reads to generate the <u>T</u>issue-level <u>A</u>tlas of <u>M</u>ouse <u>I</u>soforms (TAMI - available at https://genome.ucsc.edu/s/vollmers/TAMI) which we believe will be a valuable complement to conventional, manually curated reference genome annotations.

## Introduction

For any model organism, a high quality reference genome sequence and accompanying reference genome annotation are invaluable research resources (McGarvey et al. 2015).

This is especially true for the mouse which has been widely used as a model organism for studying basic biology and biomedical research for almost 100 years. Mice are small, easy to care for, and have short lifespans. Inbreeding of mice has led to genetically identical strains allowing for reproducible experiments. They share over 15,000 protein coding genes with humans and are susceptible to many of the same diseases (Eppig et al. 2015). Mice are easily genetically engineered to simulate many human conditions. These features combined make mice critical for scientific research.

The initial mouse reference genome was published 20 years ago and has been improved since then to be highly complete and contiguous (Mouse Genome Sequencing Consortium et al. 2002). In contrast, a truly comprehensive reference genome annotation for the mouse does not currently exist. Current reference genome annotations like GENCODE and RefSeq contain the locations of genes, their exons, and how these exons can be combined into transcript isoforms (Frankish et al. 2019; Kawai et al. 2001; McGarvey et al. 2015; ENCODE Project Consortium 2004). These reference genome annotations are absolutely essential for virtually all transcriptomics research and beyond, but they lack information on isoform composition and expression levels in different tissues. A resource containing this tissue-level isoform information would be highly useful for the design of a range of assays that require knowledge of any gene of interest in any given tissue - from the design of CRISPRi probes, RT-qPCR primers, overexpression vectors, and beyond.

In the last few years, third generation long-read sequencing technology in the form of Oxford Nanopore Technologies (ONT) and Pacific Biosciences (PacBio) sequencers and their cDNA library preparation protocols have matured. Using library preparation like MAS/Kinnex and R2C2, these sequencers are now capable of generating many millions of highly accurate sequencing reads that are thousands of nucleotides in length (Byrne et al. 2019a). For the analysis of transcriptomes this means that entire full-length transcripts can be captured as single reads, including the poly(A) tails, transcription start sites (TSS), and all splice junctions.

In theory, this type of throughput and accuracy makes it possible to generate accurate transcript isoform expression information for many tissues. In fact, ENCODE and GTEx consortia have generated deep full-length cDNA datasets for many human organs (Glinos et al. 2022; Reese et al. 2023). However, while ENCODE also generated mouse data, it did so for only a few tissues.

Here, we generated the most comprehensive genome annotation for the mouse to date. Further, by doing so in a streamlined and cost efficient way, we provide a blueprint for future genome annotation efforts of other organisms. To achieve this, we used the nanopore-based R2C2 long-read sequencing method (Volden et al. 2018; Adams et al. 2020; Vollmers et al. 2021; Cole et al. 2020; Volden and Vollmers 2022; Byrne et al. 2019b) which increases read accuracy and decreases length biases of ONT sequencers to generate over 60 million accurate full-length cDNA reads across twelve tissues from the BALB/c mouse strain. We analyzed these reads with the Mandalorion isoform identification pipeline which combines Specificity and Recall (Pardo-Palacios et al. 2022; Volden et al. 2023) to create a high quality genome annotation containing information on which isoforms are expressed at what level for each gene in each tissue. In addition to creating and releasing this <u>T</u>issue-level <u>A</u>tlas of <u>M</u>ouse <u>I</u>soforms (TAMI) - available at https://genome.ucsc.edu/s/vollmers/TAMI, we also used its underlying dataset to investigate how isoform usage varies across tissues.

## Results

### Generating accurate full-length cDNA data from 12 mouse tissues

We constructed tissue-level, long-read transcriptome data using commercially available high quality RNA (Takara) from 12 mouse tissues (brain, eye, heart, kidney, lung, liver, salivary gland, smooth muscle, spinal cord, spleen, testis) each pooled together from dozens to hundreds of BALB/c mice (Figure 1). We prepared full-length cDNA using a modified Smart-Seq2 protocol (see methods). To increase sequencing coverage of longer transcripts, which are biased against in the sequencing process, some of the cDNA was size-selected for molecules >2 kb in length by gel electrophoresis.

**Figure 1.**
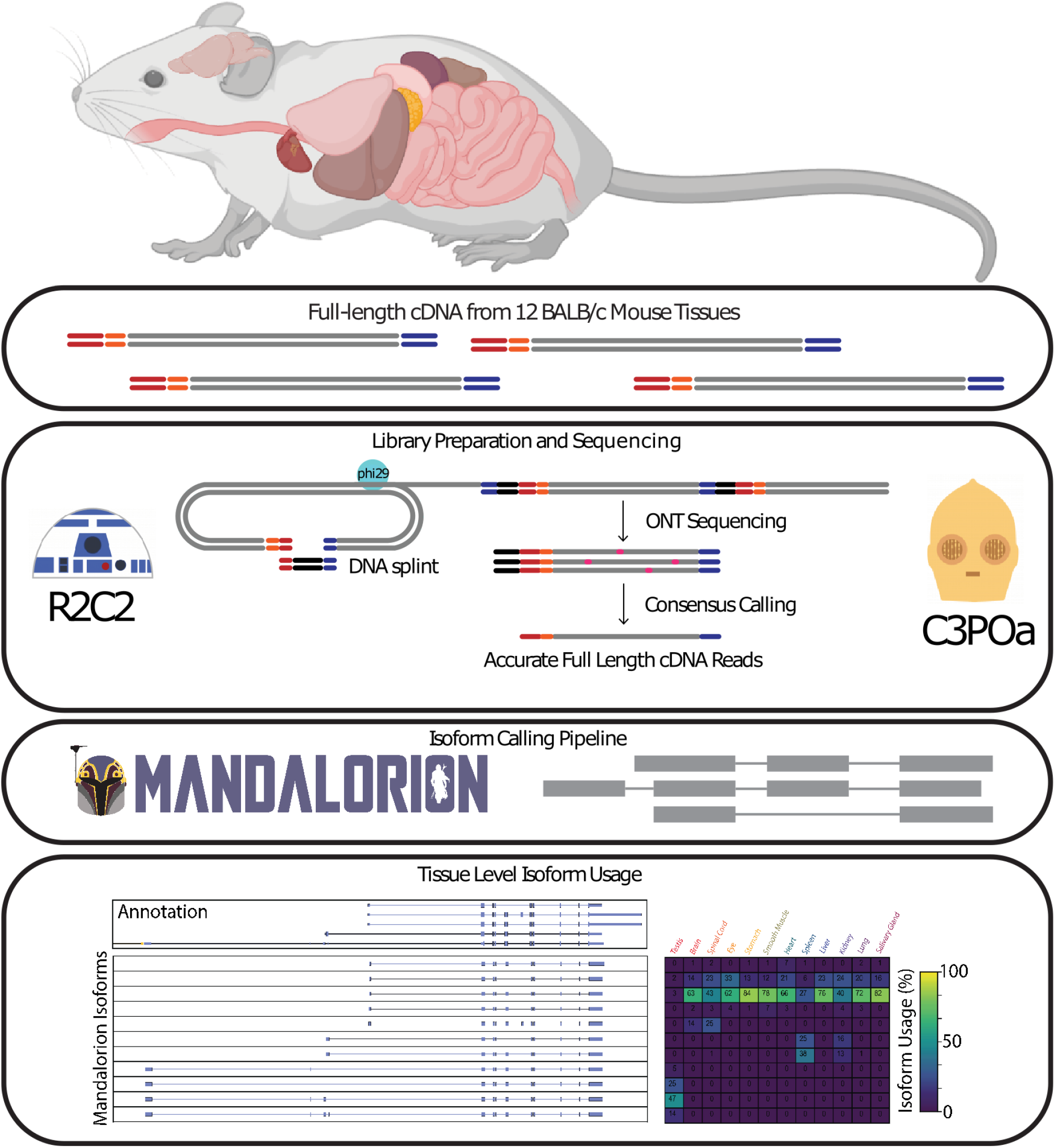
Experimental Overview. Full-length cDNA was created from total RNA extracted from 12 BALB/c mouse tissues. Pooled cDNA, both non size-selected and size-selected, was prepared for ONT sequencing by the R2C2 method. ONT raw reads were demultiplexed and consensus called using C3POa. To generate a tissue level transcriptome for each tissue R2C2 consensus reads were then processed into isoforms using the Mandalorion pipeline.

We then prepared non size-selected (*nss*) and size-selected (*ss*) full-length cDNA for ONT sequencing using the R2C2 protocol and sequenced the resulting DNA on a combination of Oxford Nanopore Technologies MinION and PromethION sequencers (R9.4 pore chemistry and SQK-LSK110 library preparation kits). After basecalling the raw signal data using Guppy (v5) we generated accurate full length cDNA consensus reads using the C3POa pipeline which also demultiplexed the resulting consensus reads into their tissue of origin. In this way, we produced 64 million full length cDNA consensus reads, averaging 5.4 million reads per tissue (Fig. 2, top). For *nss* libraries the median insert length was approximately 750 bp while the *ss* libraries had median insert length approximately 2kb (Fig 2, center). Further, the full-length R2C2 consensus reads were very accurate, with the median per base identity for *nss* and *ss* reads being 99.8% and 98.9%, respectively (Figure 2). The lower accuracy of the *ss* reads was due to longer cDNA inserts being covered less often by ONT raw reads.

**Figure 2.**
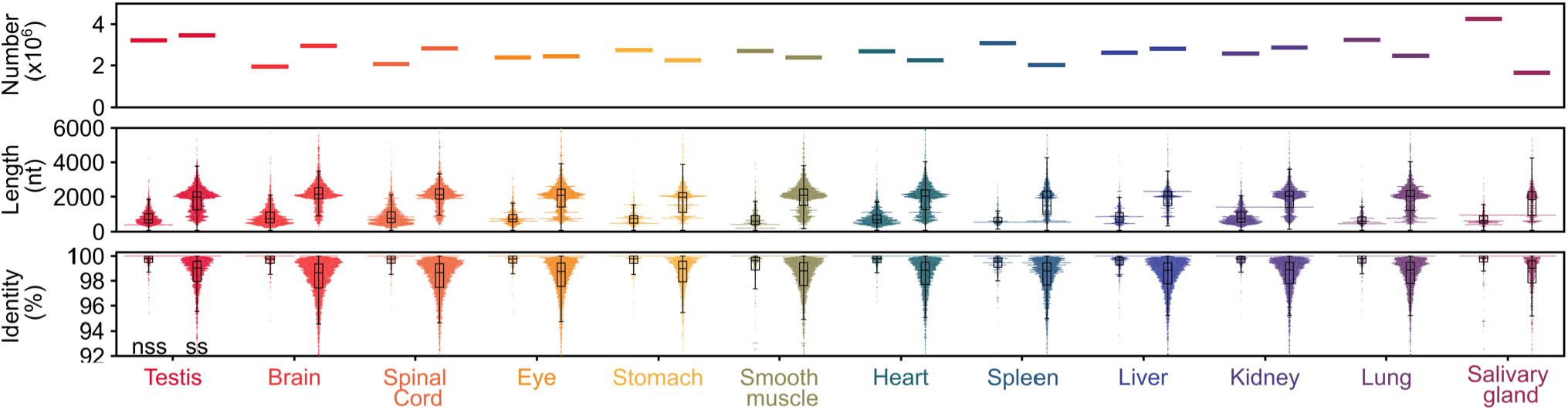
R2C2 read characteristics. Top panel, read counts in millions split between non size-selected (nss) and size selected (ss) libraries. Center panel, read accuracy of C3POa full length consensus reads split between *nss* and *ss* libraries. Bottom panel, insert length split between *nss* and *ss* libraries.

### Evaluating gene level expression quantification

Next, we determined whether the full-length cDNA R2C2 reads we generated could be used for gene detection and expression quantification. The first analysis we performed aimed to determine if our sequencing depth was enough to capture all genes expressed in the samples. We compared gene saturation of our data set (Figure 3A) to publicly available Illumina data (Figure 3B) generated for the same twelve tissues (Brawand et al. 2011; Huntley et al. 2016; Gluck et al. 2016; Merkin et al. 2012; Mustafi et al. 2011; O’Rourke et al. 2015; Söllner et al. 2017), and SRA accession SRR2927121. Comparing data sets, we saw the R2C2 data set approaching a plateau but with fewer total genes detected then the Illumina data. This can be at least partially attributed to the almost 10-fold difference in read counts.

**Figure 3.**
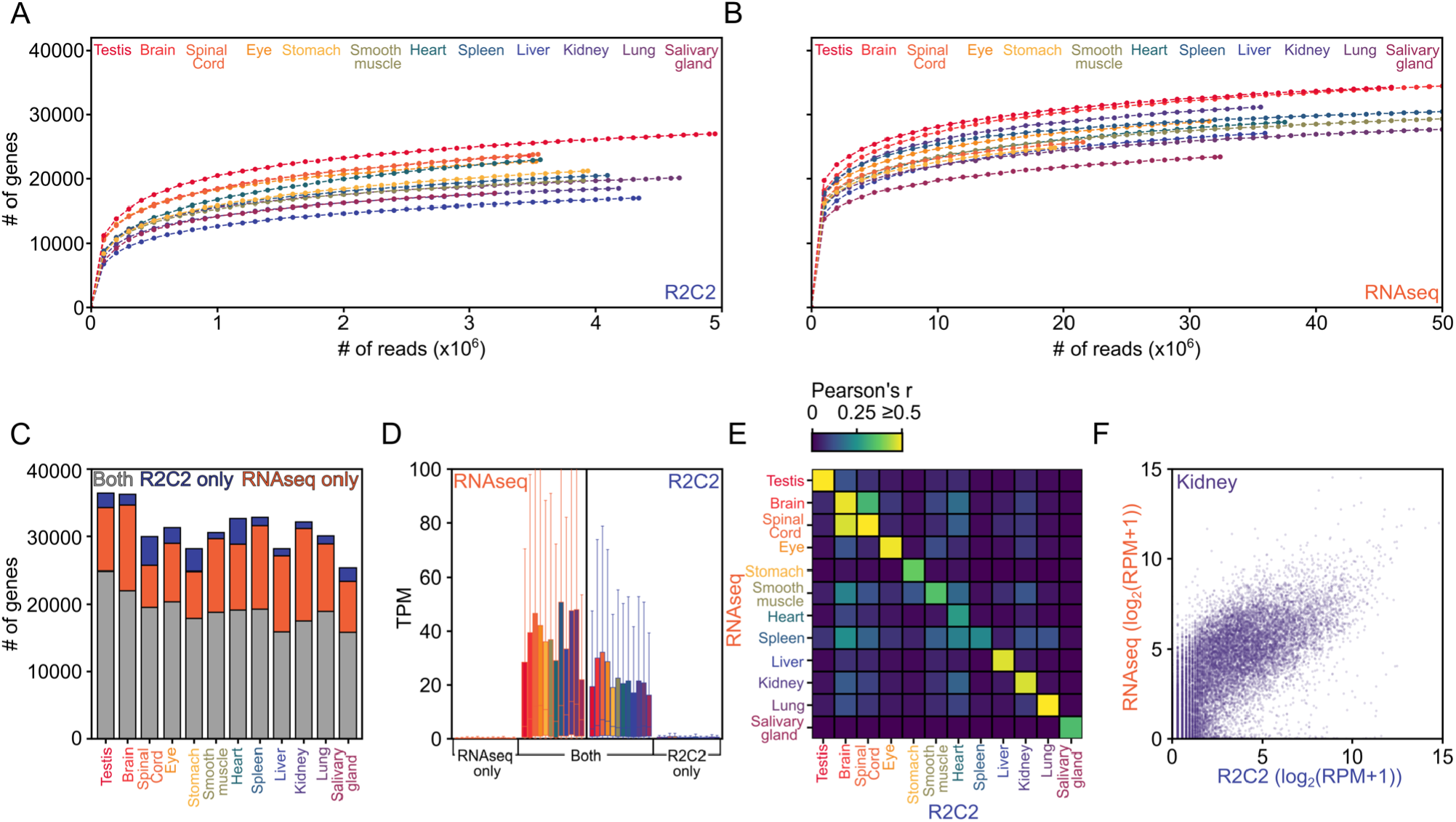
R2C2 and RNAseq detect largely the same genes at similar levels. Gene level saturation curve analysis of R2C2 data (A) and Illumina RNAseq data (B). C) Comparison of genes detected by either R2C2 or Illumina RNAseq or both. D) Expression levels as determined by R2C2 and RNAseq for genes detected by either R2C2 or Illumina RNAseq or both (Colors as in A). E) Pearson correlation between gene expression values as determined by R2C2 and RNAseq for each tissue. F) Scatterplot of kidney gene expression values as determined by R2C2 and RNAseq.

Then, we compared the number of genes detected by either R2C2 only, RNAseq only, or both methods. (Figure 3C). The majority of genes detected were identified by both methods and more genes were identified by RNAseq only then R2C2 only. However, the genes that were identified by only one of the two methods were generally expressed at very low levels (Figure 3D). This also explains why genes might be missed by either method and also why RNAseq with its higher read count might detect more genes than R2C2 (Figure 3D).

In addition to detecting genes, we analyzed whether R2C2 was quantitative in determining their expression. We did so by using featureCounts(Liao et al. 2014) to count the number of RNAseq and R2C2 reads that align to a particular annotated gene for each analyzed tissue. After converting these counts to <u>R</u>eads <u>P</u>er <u>M</u>illion (RPM) we compared R2C2-derived to RNAseq-derived gene expression for each tissue. We found that R2C2 gene expression was most correlated to RNAseq gene expression for the same tissue with pearson r value ranging from somewhat correlated (Spleen r=0.22) to well correlated (Lung r=0.78). Neuronal tissues (Brain, Spinal Cord, Eye) also showed a high correlation between tissues (Figure 4E).

**Figure 4.**
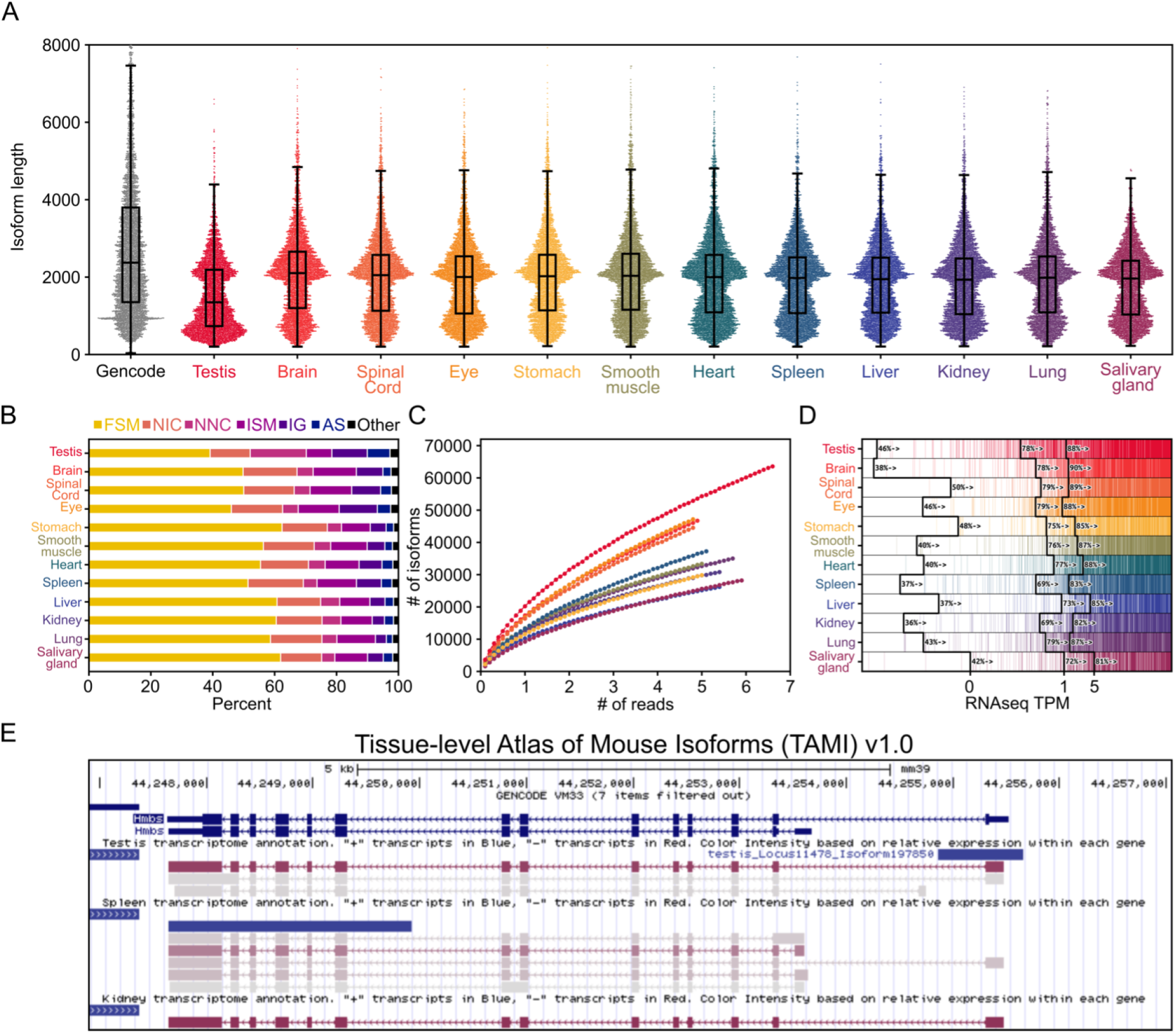
Characterization of tissue-level transcriptomes. A) Isoform length distribution for each tissue compared to GENCODE vM30 basic protein-coding transcripts. B) Isoform categories distribution for each tissue as determined by SQANTI3 C), Isoform saturation curves for each tissue. D) For each tissue, genes are rank-ordered based on their expression level in Illumina RNAseq data. Genes are marked by a vertical colored line if at least a single isoform is assigned to them in the respective tissue. 0,1, and 5 TPM levels in RNAseq data are indicated by black lines. The percentage of genes expressed higher than that TPM with at least one isoform assigned to them is shown adjacent to those lines. E) Screenshot of the testis, spleen, and kidney tracks from the TAMI trackhub as displayed on the UCSC Genome Browser.

The lower correlation between some R2C2 and RNAseq in some tissues could be due to biological differences between the RNA samples we used and those underlying the publicly available Illumina data, highlighting the limitation of using publicly available data. Generally, the gene overlap and high expression correlation between R2C2 and Illumina data in at least some tissues suggests that combining R2C2 data from *ss* and *nss* cDNA does not substantially distort gene content and expression.

### Characterizing tissue-level isoforms

To take full advantage of our long-read data, we aimed to use the full-length R2C2 consensus reads to move beyond gene level analysis and define comprehensive sets of isoforms for each of the 12 tissues in this study. To identify isoforms in a way that has high Recall and Specificity, especially with unannotated isoforms, we analyzed the R2C2 reads we produced using the Mandalorion (v4.0) tool (Volden et al. 2023). Mandalorion identifies, filters, and quantifies isoforms to create high confidence sets of transcript isoforms.

For the individual tissue data sets, Mandalorion identified between 22,727 (salivary gland) and 63,948 (testis) isoforms. Across tissues, isoforms were about 2kb in median length with the exception of testis which contained shorter isoforms overall.

To characterize the isoforms Mandalorion had identified for each tissue, we used SQANTI3 (Tardaguila et al. 2018). SQANTI3 classifies isoforms into four main categories based on comparison to the reference annotation file: full splice match (FSM), incomplete splice match (ISM), novel in catalog (NIC), novel not in catalog (NNC). For FSM, the isoform must fully match the splice junction chain of an annotated isoform but the exact 5’ and 3’ ends can differ from the annotation. For ISM, the isoform must match a partial splice junction chain of an annotated isoform but be incomplete on either 5’ or 3’ end. For NIC, the isoform must use only annotated splice junctions but in an unannotated combination. Finally, for NNC, the isoform must use at least one unannotated splice junction. Across tissues we see about 80-90% of isoforms falling into the four main categories at similar ratios, again with the exception of testis with which showed a high number of NNC isoforms indicating the use of many unannotated splice junctions (Figure 4B).

To investigate whether we sequenced these transcriptomes to exhaustion, i.e. more reads would not result in more isoforms being identified, we performed a saturation analysis for each tissue (Figure 4C). We did not reach saturation for any tissue but wanted to determine that we at least identified one isoform for each expressed gene. We found that, on average across tissues, Mandalorion identified at least one isoform for ∼75% and ∼86% of genes with greater than 1 TPM and 5 TPM expression levels in the Illumina RNAseq data, respectively (Figure 4D).

Overall, this showed that the isoforms we identified for each tissue were similar in length to protein-coding transcript isoforms present in GENCODE. However, the isoform we identified mostly lacked the long tail >6kb of isoforms present in GENCODE annotations likely due to R2C2 read length limitations. SQANTI3 analysis showed that while many of the isoforms we identified matched the GENCODE annotation (FSM), we identified tens of thousands of new, high confidence isoforms (NIC,NNC). Further, while we did not reach isoform-level saturation, we were able to capture at least one isoform for the majority of expressed genes which should make these tissue-level transcriptomes a valuable resource for researchers.

### Tissue-level Atlas of Mouse Isoforms

To make this resource as easily accessible for researchers as possible, we have created a trackhub for the UCSC Genome Browser (Navarro Gonzalez et al. 2021). Entitled Tissue-level Atlas of Mouse Isoforms (TAMI) this trackhub is available at https://genome.ucsc.edu/s/vollmers/TAMI for the mm39 version of the mouse genome. TAMI contains separate tracks for each tissue (Figure 4E) which contain isoform models identified by Mandalorion for that tissue. Isoform expression-levels are normalized within each gene and that normalized expression is then shown by the color of each isoform. Absolute expression in TPM can be seen by positioning the cursor over an isoform. An alignment between the R2C2 read-based consensus sequence of each isoform and the corresponding genomic sequence is available by clicking the isoform. These alignments might highlight potential sequencing errors as well as variation between the BALB/c isoforms and mm39 genome which is based on the C57BL/6 strain. Overall, the goal of the TAMI track is to give research fast and intuitive information about their gene of interest

### Investigating unique TSS usage in testis

When creating and inspecting the TAMI tracks for release, we observed that, often, isoforms expressed in testis would use unique, testis-only TSSs. This made sense considering testis is known to be the most transcriptionally complex tissue in mammals in terms of the number of expressed genes and isoforms (Kaessmann 2010).

Systematic analysis confirmed this unique TSS usage. We found that the 63,948 isoforms expressed in testis originated from 31,158 non-overlapping TSSs. Of those, 16,522 were unique to testis. This number of unique tissue restricted TSSs in the testis was much higher than any of the other tissues we investigated (Figure 5A).

**Figure 5.**
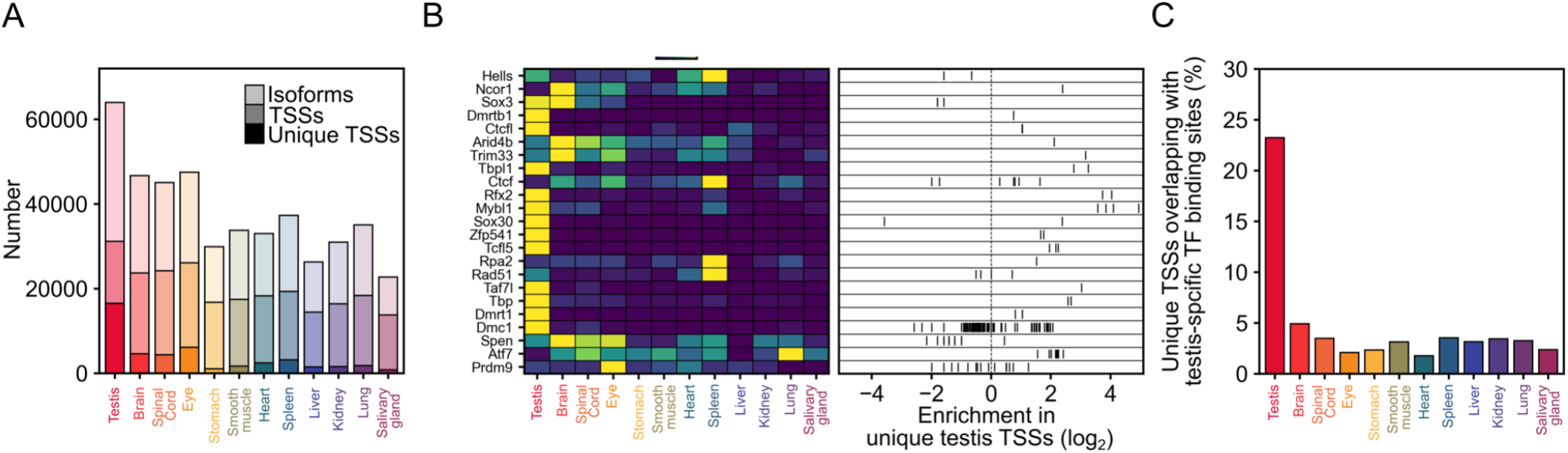
Unique transcription start sites in testis are bound by testis-specific transcription factors. A) Number of isoforms, overall TSSs. and TSSs unique to a tissue are shown for each tissue B) Left, tissue-level expression of transcription factors present in the ChIP-Atlas testis data. Right, transcription factor binding site enrichment in TSSs unique to testis. Each ChIP experiment present in the ChIP-Atlas is represented by an individual bar C) Percent overlap of the unique TSSs of each tissue with testis-specific transcription factor binding sites.

Next, we wanted to see whether the TSS unique to testis could be validated with transcription factor ChIP-Seq data from ChIP-Atlas.org (Oki et al. 2018). First, we evaluated the expression patterns of the transcription factors that had been investigated in testis (Figure 5B, left). We found that, based on our R2C2 data, several of these transcription factors were indeed most highly expressed in testis. The binding sites of these testis-specific transcription factors, as determined by many distinct ChIP experiments, were generally enriched in TSS unique to testis (Figure 5B, right).

Finally, we selected a single ChIP-seq experiment for just 6 testis-specific transcription factors - Taf7l, Tcfl5, Sox30, Mybl1, Rfx2, Tbpl1 (Yin et al. 2021; Zhou et al. 2017; Cecchini et al. 2023; Zhou et al. 2013; Kistler et al. 2015; Martianov et al. 2016). We found that their binding sites overlapped with 23% of unique testis TSSs but only ∼3% of the unique TSSs of the other tissues.

This indicated that, within the testis, testis-specific transcription factors create isoform diversity through the use of unique TSSs. The high percentage of NNC isoforms in testis suggests that those unique TSS are often unannotated.

### Differential Isoform Usage Across Tissues

The quantitative nature of the R2C2 approach as well as the multiplexed setup of our sequencing strategy allowed us to compare isoform expression across tissues. To avoid the complexity of merging 12 individual annotations, we used Mandalorion to identify isoforms from the combined data set of all 12 tissues and to quantify the expression of those isoforms in each tissue.

First, we evaluated which tissues were expressing the same isoforms - at any level - by calculating Jaccard Indexes for each pair of tissues. Again, testis proved an outlier, having lower Jaccard indexes, i.e. the smallest overlap of isoforms, than any other tissue. As expected, neuronal tissues (Brain, Spinal Cord, and Eye) had high Jaccard Indexes with each other. Stomach and smooth muscle (small intestine), both part of the digestive system, also had a high Jaccard Index.

Second, to systematically identify genes with differential isoform expression across tissues, we first identified 7,457 genes that had a combined isoform expression of at least 50 R2C2 reads (∼10TPM) in at least 2 tissues. We then performed a Chi-squared contingency table test on the relative isoform usage of each of those genes. After Bonferroni correction we found 3,742 genes with significant differential isoform usage at p≤0.01. An example of one such gene, Rab3il1 shown in Figure 6, highlights differential isoform usage across tissues particularly in regards to the use of alternative TSS and first exons, as well as alternative internal exon usage within the same tissue.

**Figure 6.**
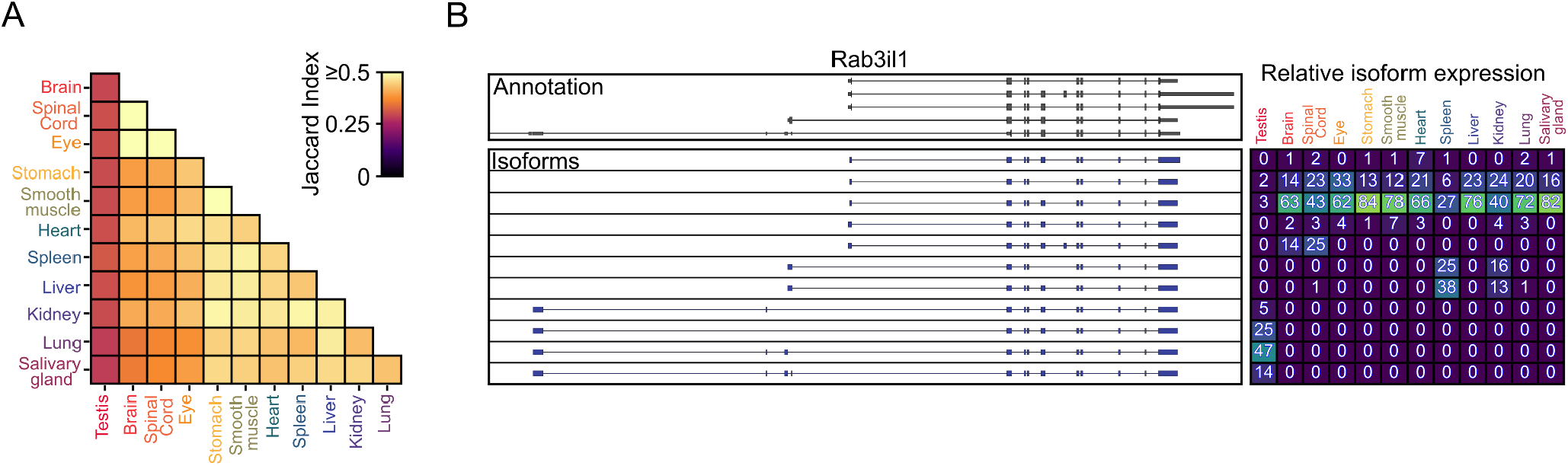
Differential Isoform Usage. A) Jaccard Indexes for each pair of tissues B) Genome Browser shot of Rab3il1 is shown with GENCODE vM30 annotation on top and isoforms called by Mandalorion below. Right side, relative usage of each isoform in each tissue, yellow indicates higher usage, blue indicates lower usage.

Overall, our analysis shows if a gene is expressed moderately high in at least two tissues, it is more likely than not (3,742 out of 7,457 or ≈50.2%) to show differential isoform expression.

## Discussion

Here, we have presented a genome annotation atlas for the mouse, highlighting the immense isoform diversity between different tissues. We used the ONT-based R2C2 method to sequence over 60 million full length transcripts across twelve tissues. We compiled these reads using the the Mandalorion tool (Pardo-Palacios et al. 2022) and the resulting isoforms formed the basis for the first release of the Tissue-level Atlas of Mouse Isoforms (TAMI) which is hosted as a trackhub on the UCSC Genome Browser.

We hope these tracks and their source files will be a valuable resource for genomics research by, for example, complementing existing annotations with tissue specific information for transcriptome dependent RNAseq analysis by tools like salmon (Patro et al. 2017) and kalisto (Bray et al. 2016). Further, by identifying more accurate transcripts ends, TAMI might improve the analysis of scRNAseq data whose reads are most often limited to the 3’ or 5’ end of transcripts.

We also hope that TAMI will be of value to bench scientists by providing easy-to-access detailed isoform information and thereby informing experimental design. For example, instead of cloning a random isoform taken from GENCODE or RefSeq for overexpression, TAMI enables you to clone the isoform that is most expressed by your gene of interest in your tissue of interest.

TAMI v1.0 represents only a starting point for our annotation efforts. We plan on using updated technologies and methods including the PacBio Kinnex method, automated cDNA size selection, and ONT R10 pores to generate more, longer, and more accurate full-length cDNA reads for a larger number of tissues and cell types. We have also recently rewritten the Mandalorion (Volden et al. 2023) tool to be capable of handling the much larger >100 million full-length cDNA read datasets that are now becoming a reality.

Finally, while TAMI is intended to merely complement the excellent manually curated reference annotations like GENCODE that are available for mouse, this paper shows that, going forward, full-length cDNA based annotation efforts could serve as reference annotations for less researched organisms.

We think TAMI provides a blueprint for these efforts. The generation of indexed cDNA makes it possible to pool samples early which in turn allows for the cost-effective generation of sequencing libraries. ONT sequencing, due to its low device cost, can be performed in most molecular biology labs. Finally, data analysis, including Mandalorion-based isoform identification, can be performed on consumer-grade computers. All of this puts genome annotation efforts within reach of individual labs with moderate budgets.

## Methods

### Sample Multiplexing

RNA was acquired from Takara (Cat# 636644). Multiplexing samples was done using one of two methods: the first used barcoded DNA splints for Gibson assembly then pooling samples after rolling circle amplification, the second method used barcoded oligo(dT) for cDNA synthesis wich allowed pooling before Gibson assembly. Both methods produce equivalent data. Approximately 80% of the data used in this study was generated by using barcoded oligo(dT) primers for multiplexing tissues.

### Library Preparation and Sequencing

#### cDNA synthesis

RNA was first mixed with dNTPs and oligo(dT) primer, either barcoded or non-barcoded, then denatured to remove secondary structure for 3 minutes at 72C. First strand reverse transcription (RT) using Smartscribe Reverse Transcriptase (Clontech) and SmartSeq template switching oligo (TSO) with DTT and SUPERaseIN was performed for 1 hour at 42C then heat inactivated for 5 minutes at 70C. Second strand synthesis and PCR with KAPA 2x master mix and ISPCR primer with RNaseA and lambda exonuclease for 12 cycles3 (37C for 30 minutes, 95C for 3 minutes, 98C for 20 seconds, 67C for 15 seconds, 72C for 8 minutes, 72C for 5 minutes, 4C hold). cDNA was cleaned up and size-selected using SPRI beads at a 1:0.85 (sample:beads). After quantification by Qubit the cDNA libraries were pooled together if barcoded oligo(dT) primers were used, if not, cDNA from individual tissues would still be kept separate. The cDNA was then split for size-selected and non size-selected R2C2 library preparation. For size selection, cDNA was run on a 1% low melt agarose gel and everything over 2 kb was excised and purified using beta-Agarase digestion and SPRI bead clean up.

#### R2C2 library generation

Size-selected and non size-selected cDNA were further processed separately but identically. cDNA libraries were circularized by gibson assembly (NEBuilder HiFi) with a short DNA split that overlaps with the ends of the cDNA. For cDNA that was not barcoded during cDNA synthesis a barcoded DNA split was used. To remove un-circularized molecules, a linear exonuclease digestion with ExoI, ExoII, and Lambda Exonuclease (all NEB) was carried out for 16 hours at 37C then heat inactivated for 20 minutes at 80C. The reaction was then cleaned using SPRI beads at a 1:0.85. The clean, circularized library is then used as a template for rolling circle amplification (RCA) using Phi29 (NEB) with a random hexamer primer for 18 hours at 30C then heat inactivated for 10 minutes at 65C. The phi29 reaction was then debranched using T7 endonuclease for 2 hours at 37C before being cleaned and concentrated using Zymo DNA clean and concentrator column. The library was quantified by Qubit and gel extracted as described above but the region extracted was a bright band just over the 10 kb marker. After gel extraction, the library was quantified again by Qubit.

#### ONT sequencing

Libraries barcoded during the Gibson assembly step were now pooled together at equal mass. We used the Genomic DNA by Ligation (SQK-LS110) kit from ONT to prepare for sequencing following the manufacturer’s protocol. The final library was loaded onto either a ONT MinION or PromethION sequencer. Flowcells were nuclease flushed and loaded with additional library partway through sequencing based on pore availability statistics shown in the MinKNOW software to increase sequencing throughput.

### Data Processing

All ONT fast5 files were basecalled using Guppy (v.5) with the super accurate configuration. R2C2 full length consensus reads were generated and demultiplexed by C3POa (v2.4.0). Isoforms were called by the Mandalorion Isoform analysis pipeline (v4.0) run on both individual tissue data and the combined data set.

### Analysis

Read identity was determined by aligning reads to the GRCm39 version of the mouse genome and identifying mismatches and indels. Gene level saturation curves were produced by random subsampling of featureCounts output for each tissue and the combined dataset. Isoform level saturation curves were produced by random subsampling FASTA files output by C3POa and running Mandalorion on each subsample.

Pearson correlation and scatter plots comparing Illumina and R2C2 gene quantification were produced using gene read counts from featureCounts. R2C2 data was aligned to the reference genome using minimap2 (Li 2018) while Illumina was aligned using STAR aligner (Dobin et al. 2013). Mandalorion isoforms produced from both individual tissues and the combined dataset were used as input for the sqanti_qc.py script of SQANTI3 v5.1 (Pardo-Palacios et al. 2023).

Differential isoform usage analysis was performed using Chi2 contingency test with a custom python script utilizing SciPy (Pardo-Palacios et al. 2023; Jones et al.; Harris et al. 2020).

A tissue unique TSS was determined by combining nearby TSS from each tissue and then comparing those to the combined TSSs of all other tissues. A tissue unique TSS was defined as a TSS in one tissue that overlaps with TSSs in any other tissue. A custom python script was used to determine TSS overlaps with publicly available ChIP-seq (https://chip-atlas.org/). Data from chip-atlas.org was downloaded as a BED file from the peak browser tool by selecting the following options: Assembly: M. musculus mm10, experiment type: ChIP TF, Cell Type Class: Gonads, Threshold for Significance: 50, ChIP Antigen: all, Cell Type: testis.

## Acknowledgements

The ONT Promethion sequencing was carried out by the DNA Technologies and Expression Analysis Core at the UC Davis Genome Center, supported by NIH Shared Instrumentation Grant 1S10OD010786-01.

## Data Availability

The full length consensus reads generated in this study have been submitted to the NCBI Sequence Read Archive under accession number PRJNA971991. Custom UCSC genome browser session with isoforms called by Mandalorion and relative usage can be found at https://genome.ucsc.edu/s/vollmers/TAMI. Isoform models in GTF format can also be found at https://vollmerslab.sites.ucsc.edu/tami/

